# Ubiquitous abundance distribution of non-dominant plankton across the world’s ocean

**DOI:** 10.1101/269068

**Authors:** Enrico Ser-Giacomi, Lucie Zinger, Shruti Malviya, Colomban De Vargas, Eric Karsenti, Chris Bowler, Silvia De Monte

## Abstract

Species Abundance Distributions (SADs) bear the imprint of ecological processes that shape biological communities, and are therefore used to discriminate among different scenarios of community assembly. Even though empirical distributions appear to follow a handful of qualitative laws, it is still unclear if and how quantitative variation in SADs reflects peculiar features of the communities and their environmental context. Here, we use the extensive dataset generated by the *Tara* Oceans expedition for marine microbial eukaryotes (protists) and an adaptive algorithm to explore how SADs vary across plankton communities in the global ocean. We show that the decay in abundance of non-dominant OTUs, comprising over 99% of local richness, is commonly governed by a power-law. The power-law exponent varies by less than 10% across locations and shows no biogeographical signature, but is weakly modulated by cell size. Our findings suggest that large-scale ubiquitous ecological processes govern the assembly of non-dominant plankton throughout the global ocean.

## Main text

Marine plankton communities are composed of an enormous number of cohabiting species ^1 2 3^, in spite of the potentially strong selective pressures imposed by the abiotic environment^4^. Plankton blooms at temperate latitudes - where a few dominant species become hugely abundant in just a few days - spectacularly illustrate the existence of spatio-temporally restrained niches^5^. Heterogeneity in the physico-chemical environment thus affects local species composition and underpins plankton biogeography^6^.

Appropriate adaptation to idiosyncratic environmental conditions and competitive exclusion are however mitigated by the strong spatio-temporal variability of ocean dynamics^7^. Therefore, a fraction of plankton communities is found in sub-optimal growth conditions, with a resulting low abundance. In microbial assemblages, this non-dominant component is referred to as ‘the rare biosphere’^8 9 10 11 12^, and is largely responsible for their enormous diversity. Since the species composing the rare biosphere are for the most part unculturable and lack precise morphological and ecological characterization, the investigation of plankton communities and their ecological patterns relies more and more on the measure of their genetic diversity. On the one hand, these observations have revealed plankton biogeography, showing that local species composition reflects the distribution of environmental drivers^13 1^. On the other hand, plankton assemblages have been included in the rapidly growing number of microbial communities used as benchmarks for testing general macroecological theories and supporting the existence of universal biodiversity patterns ^14 15 8 16^.

One primary feature of ecological communities, on which classical indicators of diversity and evenness are based, is how many individuals belong to each of the taxa observed in a local sample^7^. This information is typically represented in two forms. In rank-abundance plots, species are arranged in decreasing order of abundance, whereas Species Abundance Distributions (SADs) represent the frequency histogram of species relative abundances. The shape of these plots is invariably characterized by a small number of abundant species and a larger number of rare ones, motivating the search for general principles able to account for the functional form of the abundance decay^17 18 19 20 21^. Attempts of deriving the shape of SADs from ecological mechanisms or statistical laws has had to address the problem of the deviations in empirical distributions from the theoretically-postulated laws. Through the development of increasingly sophisticated statistical methods^22 23 24^, the resemblance between empirical and theoretical distributions has been assessed quantitatively. This had raised the hope of leveraging the increasing data from diverse communities - heterogeneous in organisms type, physical location, and sampling method - to identify universal first principles able to encompass all observations within a unique framework^22 23 25^. The results of such efforts are however far from clear. Even though the log-normal (or Poisson-lognormal) and log-series models appear to have the best performance overall^22 23 26^, the best-fitting functional form not only varies from community to community, but seemingly depends on the environmental context, the biome, and the sampling effort^22^. One possible solution to the failure of demonstrating universality based only on fitting SADs is to try to fit other ecological patterns, such as the variation of taxonomic abundances in space and time^27^ and body-size distribution^26^, in order to increase the discriminating power of the models. However, these extensions require a spatiotemporal and/or morphological resolution in community analyses that is seldom available in molecular datasets.

The identification of universal laws by fitting empirical distributions with one of the many available theoretical models may also be hampered by limitations of model selection approaches. First, by focusing on the best-fitting theoretical distribution, these approaches do not examine the parameters of the best fit. Since different models can provide very similar distributions (for instance, the Poisson-lognormal can yield a power-law behaviour) and models based on distinct assumptions predict, for appropriate parameter values, effectively indistinguishable distribution^26^, different families of laws can fit equally well an empirical distribution. Furthermore, it is usually assumed that the ecological processes or statistical principles underpinning empirical distributions apply uniformly to all taxa in the community. Whereas this might hold true for theories based on system-level constraints, it would be natural to expect that different components of the community are best described by different process-based models, such that empirical SADs are the overlap of two or more theoretical distributions. Magurran & Henderson^28^ and Ulrich & Ollik^29^ have indeed shown that ‘occasional’ and ‘frequent’ (in a time series) species in animal communities follow different SADs, with the former obeying a highly skewed distribution. Fitting the whole community with a single law would in this case introduce irreducible discrepancies, whose magnitude would moreover vary from one location to the other, as the proportion of taxa in different components of the community changes.
Here, we take a novel approach that allows us to examine the quantitative variation in the distribution that best fits the empirical SADs across communities that are ecologically uniform (marine plankton), but differ both in taxonomic composition and in physical location in the global ocean. We restrict our focus to taxa that are non-dominant, and which in communities subjected to a high degree of dispersion are arguably akin to ‘occasional’ species. In order to reduce the possible bias introduced by choosing a specific family of fitting distributions, we use several laws considered in the literature, including a general functional form that encompasses generic shapes of abundance decay (exponential and power-law). Our analysis of the distributions fitting best the rare component of the community reveal an unexpected quantitative regularity in their parameters, which we interpret as the signature of shared ecological processes that make the ecology of non-dominant taxa effectively neutral.

### Results

We exploited the extensive molecular dataset generated by the *Tara* Oceans expedition, which comprises millions of protist metabarcode sequences from 121 different locations worldwide^30^, constituting a comprehensive picture of planktonic protist communities. By using different proxies for protist ‘taxa’, as defined by swarm^31^ (hereafter swarms or species) and UCLUST for 95% and 97% sequence identities (hereafter OTUs), we aimed to minimize possible intraspecific variability and PCR/sequencing errors. We analyzed abundances of pelagic protist taxa spanning more than three orders of magnitude in cell size (from 0.8 μm - 2 mm, subdivided in 4 size classes: pico-nano, nano, micro, meso; Methods), hence assessing communities that are subjected, other than to environmental heterogeneity, to different physiological and ecological constraints in terms of their metabolic rates, population densities, dispersal and diversification potential^32 33 34^. The depth of sequencing^2^ and consistency of sampling protocols (Methods) allowed us to address not only the way relative species abundance decays in every local community, but also its geographical variation^35^.

As expected, considering that they are sampled in different locations and environmental contexts, the overlap in composition of any couple of the 121 communities examined was small. The average Jaccard index is 0.14 ± 0.07 for the pico-nano size class, 0.08 ± 0.06 for nano, 0.08 ± 0.07 for micro and 0.03 ± 0.06 for meso. Correspondingly, the rank-abundance plots of planktonic protists displayed sample-to-sample heterogeneity, as illustrated in Fig. 1a. However, the tail of non-dominant species - responsible for most of the community diversity - appears to follow an analogous scaling in all samples. This regularity may indicate that species in the range of lowest abundances can be considered separately from the most abundant plankton types. If local communities are the composition of groups of species that are driven by distinct ecological and/or statistical processes, we can try to isolate a single group based on its adherence to the same scaling law. We thus decided to use the homogeneity of the abundance decay as a proxy for automatically identifying, sample-by-sample, the community component of non-dominant taxa. We adapted the method introduced in Clauset et al.^36^ (Methods) to single out the largest community component that is described by a family of distributions within a given level of statistical confidence (Methods). We applied such an adaptive algorithm to several theoretical models, including classical distributions^18 21 37^ (exponential, log-series, log-normal, Poisson log-normal and zero-sum multinomial) and to a general model, described below, with density-dependent species-neutral demography, which includes both exponential and power-law decay^38 39^. Models having a probability lower than 10% of generating a distance equal or larger than the observed distance between empirical and best fitting distributions were rejected as being not statistically significant (corresponding to p<0.1; see Methods). The largest range of abundance of statistically significant fits (the extension of the fit) singles out the non-dominant component of the community that is comprehensively described by the chosen family of distributions. We refer to the distribution that has the maximum likelihood over the extension as the best-fitting distribution of a given sample (e.g., in Fig. 1b). A definition of the rare component based on setting an arbitrary (and sufficiently small) threshold in abundance^13^ would provide similar best-fit parameters, but underestimate the extension of the non-dominant component, thus also decreasing the statistical power of the fit. Importantly, since fitting distributions belong to the same functional family, the parameters of the best fit are quantitatively comparable across organism size and geographical location.

**Figure 1.**
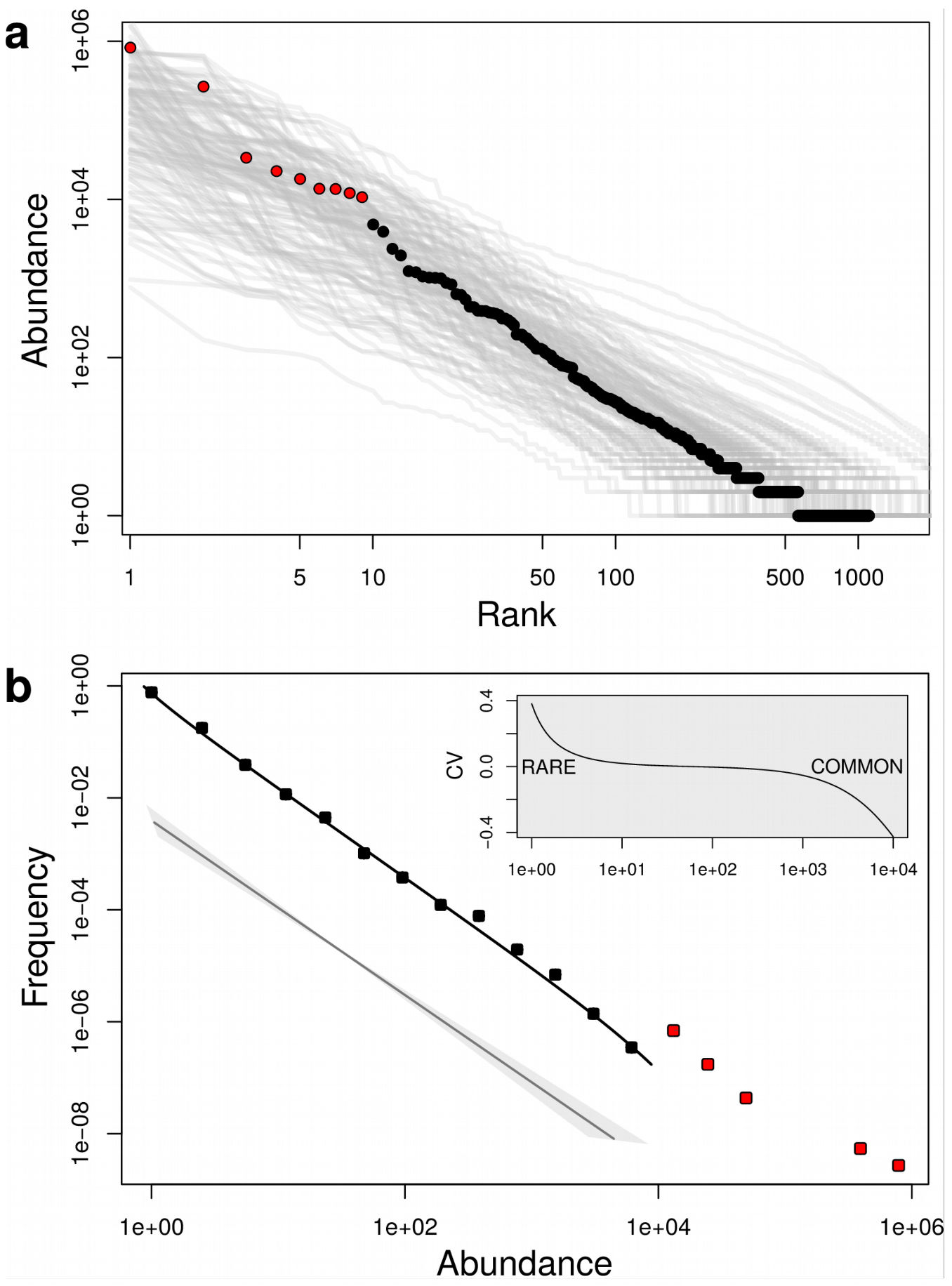
Abundance Distributions of Protist Swarms. **a,** rank-abundance plots for all local communities (grey) reveal that the abundance decay for non-dominant species is consistent across samples. Circles illustrate how dominant and non-dominant species get partitioned in one local community (same colours as in panel b). **b,** The corresponding empirical SAD (squares) is the histogram of swarms of similar abundance (logarithmic binning). The frequency of non-dominant species displays a different regime (black), than dominant ones (red). The distribution obtained by adaptively fitting the model equation (3) (black curve), describing the non-dominant component of the community, has a leading associated power-law trend whose exponent λ is given by equation (1). The limited range of variability across locations and sizes of such trend is illustrated below in grey. The straight gray line is a power-law with exponent corresponding to the average of λ across all samples. The gray shaded region represents the extent of the power-law variation when the exponent varies within one standard deviation around such mean value. The inset displays the coefficient of variation between the best-fitting distribution given by equation (3) and its associated power-law, showing that the model can capture deviations from the power-law scaling for both abundant and rare species. The distribution used for this illustration has been chosen among those which display deviations from a power-law regime for both low and high abundances, and have a sizable dominant component. The fits for all samples are displayed in Supplementary Data 1.

Examination of the empirical abundance distributions indicates a dominant power-law trend in the abundance decay, with no multimodality. We hence chose to fit the data with a model that reproduces this distribution and can moreover accommodate deviations from the power-law regime, which are common for the rarest abundance classes of empirical SADs (Methods). This model (equation (3), Methods) has been derived by He^38^ under the neutral hypothesis that demographic events (birth and death) occur for every species at the same constant per capita rates *b* and *d*, corrected by nonlinear terms χ and μ^38 27 39^ that depend on the number of individuals of the focal species. Such nonlinearities can be interpreted as describing frequency-dependent demography and/or immigration/emigration. By deriving analytically the asymptotic limit of equation (3) (Supplementary Information) we find that the model distribution encompasses three regimes (illustrated in Fig. 1b inset). Intermediate abundances decay as a power-law with exponent λ, given by:

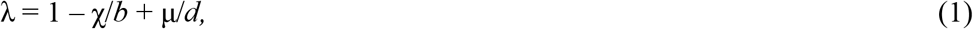

that crosses over to an exponential decay (with exponent *r*=−log(*b*/*d*)) for large abundances. On the other hand, nonlinearities in demographic rates parametrize the finite-size deviation from the power-law, that occurs for rare species and, at first order, is proportional to χ/*b* + μ/*d*. Both the exponent λ of the associated power-law and the rare species deviations thus depend on the two parameters χ/*b* and μ/*d*, which are independently fitted to the data. Equation (1) predicts that whenever the difference between the two nonlinearity parameters is constant (therefore, χ/*b* = const + μ/*d*), the power-law exponent will be invariant, independently of how rare classes deviate from the power-law trend.

The model distribution (equation (3), Methods) fits the non-dominant component of the swarm-based SADs in 372 out of 388 samples (Methods, see Supplementary Data 1 for SADs and fits of each sample). The fits to other distributions, delivered by the adaptive algorithm with the same stringent thresholds for tolerance and fit extension, were only significant in a few samples, if at all. The Poisson-lognormal distribution provides a statistically significant test for 136 samples, whereas the exponential, log-series, log-normal, and zero-sum multinomial distributions are never significant. The 16 samples that were not fitted by the model of equation (3) possessed highly irregular trends that, at the same level of statistical significance, were not fitted by any other examined distribution either. We also tested whether the fitted parameters were affected by sequencing depth (i.e., number of reads obtained for each sample), as is common in community-level statistics^18^, and found no significant effect (Supplementary Information). The possibility of fitting almost all samples with the same model allowed us to compare the parameters of the best fit across geographical sites. Table 1 reports the average and Coefficient of Variation (subdivided by size class and all size classes confounded) of the parameters for all samples where the fit was significant.

**Table 1.**
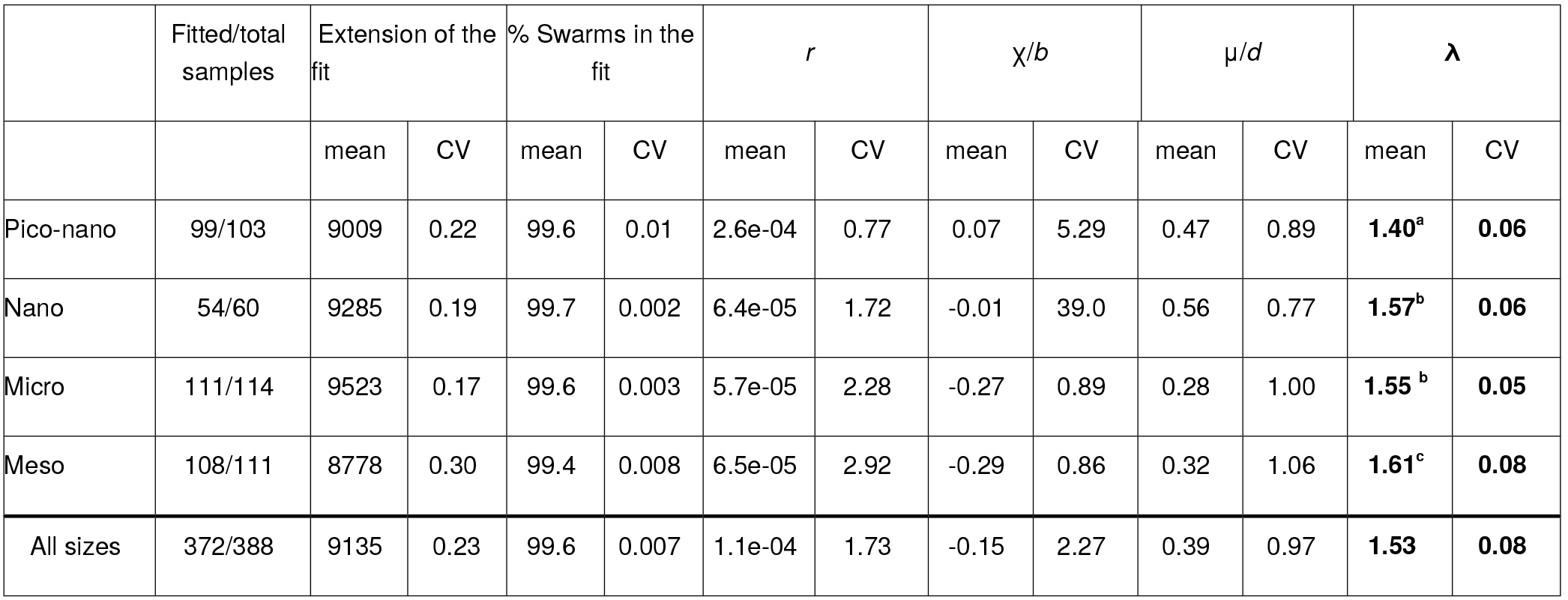
Parameters of the best fitting model. Results of the adaptive fit of the swarm-based SADs with equation (8) (Methods) and corresponding parameter statistics (mean and coefficients of variation across samples), separated by size fraction (pico-nano 0.8-5 μm, nano 5-20 μm, micro 20-180 μm, meso 180-2000 μm). The vast majority of samples can be fitted by the model to the confidence level of 0.1. The extension of the fit indicates that the model is followed for nearly 4 orders of magnitude in abundance, encompassing a broad nondominant community. Despite variations in sampling location of the fit parameters (r, χ/*b*, μ/*d*), the associated power-law exponent λ has a coefficient of variation (< 8%) much smaller than any other contextual parameter (Methods), suggesting that rareness in plankton communities follows a ubiquitous scaling law. Superscripts behind λ values indicate groups assigned by the post-hoc Tukey’s test (corrected p<0.05) for differences between size fractions.

On average, the fit encompasses 99.4% of species in each sample, and only fails to account for the abundance of a handful of dominant swarms. For non-dominant species, on the other hand, the rate of exponential decay *r* is very small throughout (Table 1). In the model, this occurs when the linear per capita birth and death rates *b* and *d* are comparable locally (though not necessarily identical in all locations), suggesting that the non-dominant component of the community is overall in demographic equilibrium. In the open ocean, such balance could be favoured by the strong dispersal induced by water transport, that would effectively temper the differences between demographic parameters (see Methods and Discussion). In the absence of an appreciable exponential cutoff, plankton protist SADs thus display an extensive power-law regime covering the largest part of the abundance range (up to 4 orders of magnitude). Moreover, the rarest classes generally exhibit a deviation (negative more often than positive) from a perfect power-law, as captured by the nonlinearity parameters χ/*b* and μ/*d*. The range of variation of these parameters is broader than what was found when fitting similar but more parsimonious models to macroorganism communities^19 39 40^. Nevertheless, they present a strong correlation along a line with slope close to 1 (Fig. 2). Such a pattern results in a strikingly small variation of the exponent of the power-law trend (equation (1)) across samples (Fig. 1b, Table 1).

**Figure 2.**
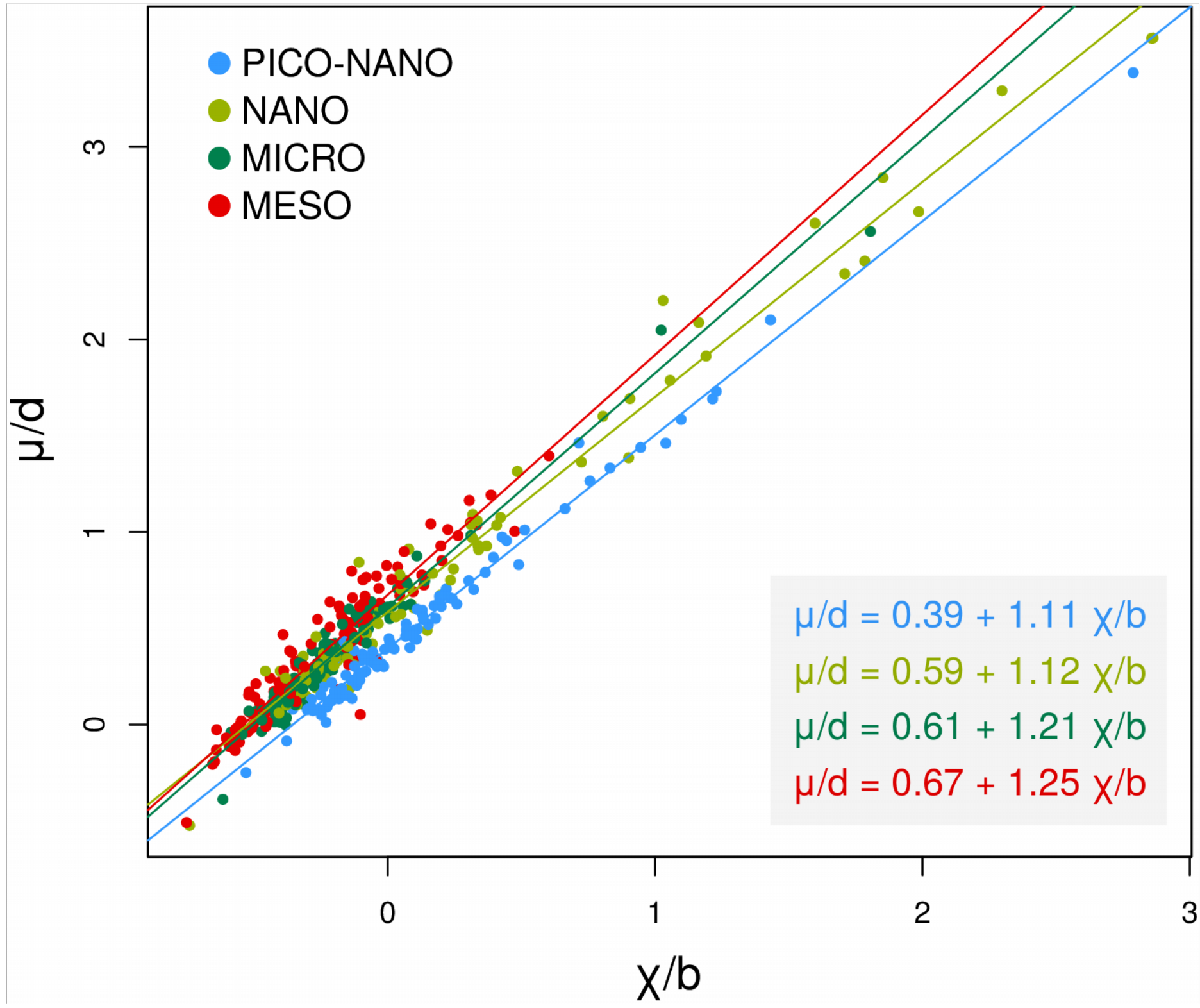
Relationship between fitted nonlinear parameters. The independently fitted relative nonlinear rates μ/d and χ/b show a very strong linear covariation, best fitted by the equations in the inset. Such linear relations (solid lines) underpin the invariance of the power-law exponent (equation (1)) within a size class: the difference of the nonlinearity parameters is almost constant even though their sum, thus the deviation from the perfect power-law regime for low abundances, has a high variability. The decrease in the slope of the regression line with size class corresponds moreover to the size-dependence of the power-law exponent, and illustrates that small size classes seem to follow more closely the relation among parameters (a line with unitary slope) predicted when the exponent λ is constant.

In terms of species composition, the dominant and non-dominant taxa of the community do not appear to be conserved across locations, as one would expect if there were species that are always rare or always common. Indeed, for each swarm we calculated the fraction of samples in which it is dominant (if present). On average, such fraction is 0.08 ± 0.15 for the pico-nano size class, 0.08 ± 0.11 for nano, 0. 09 ± 0.09 for micro and 0.07 ± 0.06 for meso. Hence, there is almost no swarm that occurs systematically in the dominant component, but swarms are instead commonly found in both dominant and non-dominant components, indicating that species that are locally abundant are in most cases rare if not absent in other locations.

When displayed on a map (Fig. 3 and Supplementary Figure 1), the uniformity of the power-law exponent value pairs with a lack of geographic signatures. We then used a collection of contextual data^41^ - including physicochemical measures performed at each sampling site, satellite-derived diagnostics, and outputs of simulations of global-scale coupled climatic and ecological models - to asses covariation with the best fit parameters^35^. Previously, local environmental variables have been shown to explain part of the bacterioplankton community features and their biogeography, and temperature is known to underpin classical macroecological patterns such as latitudinal diversity gradients^42^. Oceanographic parameters related to the processes of transport and mixing at the local scale are moreover expected to influence patterns of dispersal and niche arrangement, thus affecting community assembly. Although such abiotic and biotic contextual variables have considerably larger coefficients of geographic variation (CV) with respect to the power-law exponent (e.g., 30% for a particularly stable physical parameter, Mixed Layer Depth Temperature; 121% for nitrates; 50% for surface chlorophyll), we found no relationship holding consistently across size classes or across parameters. The few significant correlations (listed in Supplementary Table 1) explained only a small fraction of the variance (Supplementary Information). This result again supports the conclusion that no specific local condition shapes the SAD of non-dominant species.

**Figure 3.**
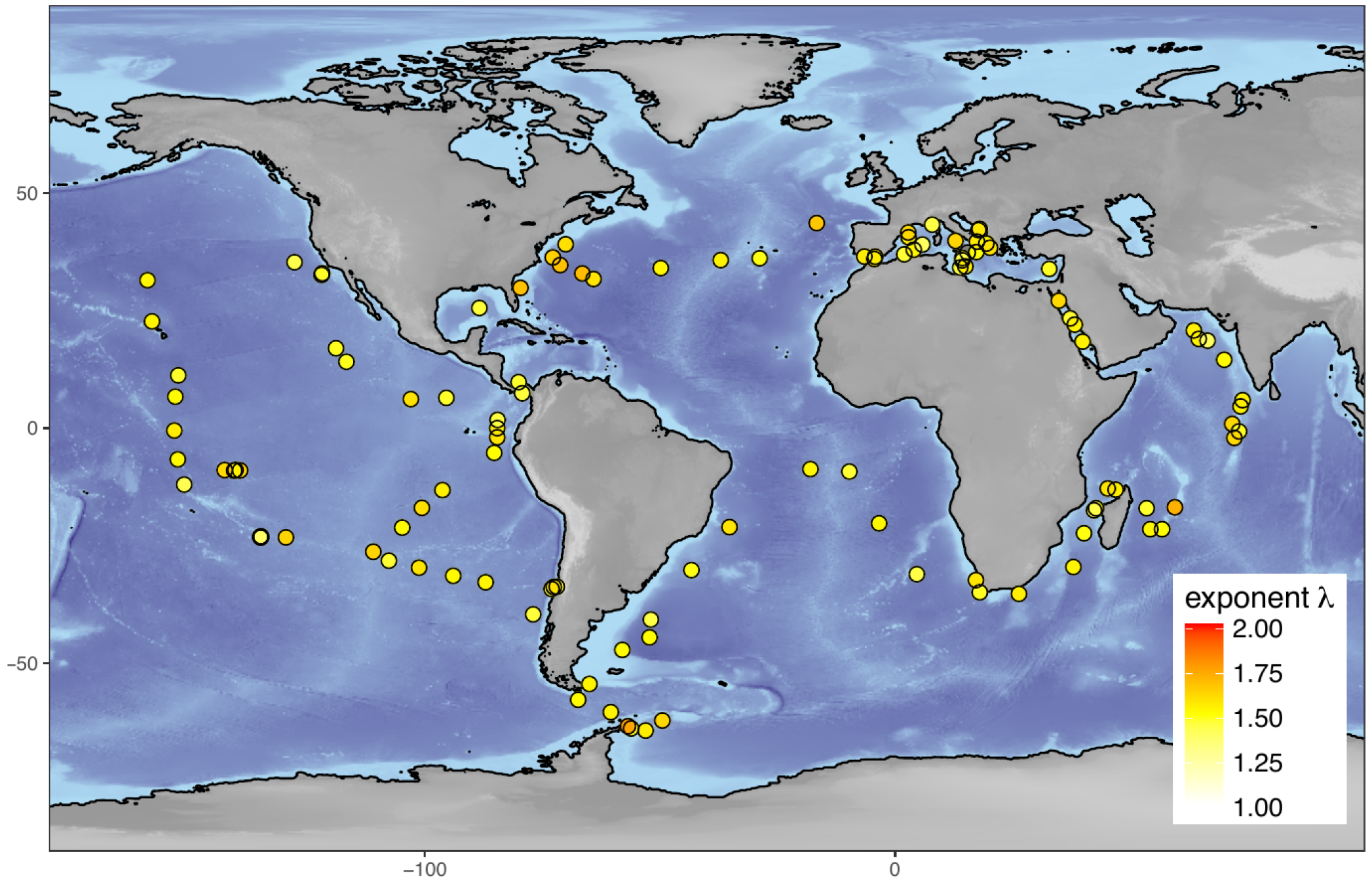
Sample locations and their associated average power-law exponent. Global distribution of the power-law exponents associated to the best fitting distributions, averaged over size classes. The color of the dots indicates the value of such exponent in a given sampling location, and displays a remarkably small geographic variation.

Consequently, dominant species are expected to carry the most biogeographical information. Indeed, those species that achieve dominance can be considered to be those that win the competition race whose terms are set by the local abiotic niche. The repartition obtained by our algorithm allows us to test if biogeographical differences in community composition can be recapitulated by considering only the most abundant species. To this end, we computed pairwise distances of communities of the same size fraction, making use of Jaccard and Bray-Curtis metrics. We then evaluated the same distances by keeping only those few ‘dominant’ species whose abundance was superior to either the threshold provided by the fitting algorithm, or to the maximal abundance we were able to process numerically (this definition of dominance is hence conservative). As illustrated in Supplementary Figure 2 and 3, these two metrics correlate very strongly independently of how the dissimilarity is measured, hence confirming the above expectations.

In spite of the overall small variation of the exponent across samples, statistically significant differences were found among size classes, with bigger size classes having the largest power-law exponent (Table 1). Systematic variations in parameters with the size class could stem from scalings in physiology, demography, or speciation rate, as the metabolic theory of ecology would predict^32^, or to size class-dependent water filtration modulating the scale of the sampled community. Alternatively, it is possible that these changes reflect the different degrees to which a neutral model is able to account for the actual process of community assembly. Due to the higher likelihood of niche overlap, it has indeed been suggested that communities with high species richness are more likely to obey the tenets of neutral theories^43^. This explanation is consistent with the observation that when size decreases (and correspondingly, richness increases), the agreement of the nonlinearity parameters to the scaling defined by equation (1) improves (Fig. 2). Moreover, plotting the exponent λ against swarm richness (Supplementary Figure 4) suggests a possible convergence of the power-law exponent to a theoretical value of 1.5 in samples displaying the highest swarm richness values.

Similar widespread power-law regimes with limited geographic variation in their exponents could be retrieved when the same analysis was repeated with OTUs (both 95% and 97%) rather than swarms.

The most notable difference is an expected dependence of the power-law exponent on the level of metabarcode sequence aggregation: its average value increases slightly but significantly as the taxonomic resolution decreases (Supplementary Table 2 and 3), indicating that abundances decay at a slower rate (and diversity increases) when the community is described at a finer taxonomic level. We therefore conclude that the consistency of the observed patterns is unlikely to be a consequence of molecular/clustering artifacts.

### Discussion

Our results demonstrate that the abundance distributions of non-dominant plankton are overwhelmingly dominated by a power-law decay. Analogies with previous work^28 29^ suggest that the non-dominant component of the community could be composed of ‘transient’ or ‘occasional’ taxa. For plankton assemblages that live in an environment with strong spatio-temporal variation and a high degree of transport-induced mixing, however, it is not obvious to distinguish endemic and occasional component. For occasional species, we can nevertheless argue that dispersal, and not competition at the sampling location, is likely to be the main driver of the demographic dynamics^44^.

By separating each community into a dominant and non-dominant part, we have found that the biogeographical information is essentially carried by dominant species, likely the most adapted to idiosyncratic environmental conditions. The non-dominant component instead follows a power-law abundance decay whose exponent varies much less than any other locally measured parameter, and that appears to carry no geographic signature.

The empirical distributions are well described by a density-/dispersal-dependent neutral demographical model, which predicts invariance of the power-law exponent when the nonlinearities in birth and death rates co-vary. The ability of a theoretical law to reproduce the empirical distributions, however, does not in itself prove that the mechanistic processes described by that model are the primary factors shaping the community. It is indeed notoriously known that the same functional form can be obtained starting from very different ecological and statistical hypotheses^17^. Nevertheless, our results suggest that distinct protist communities are fundamentally shaped in ways that are not only qualitatively the same, but lead to quantitatively similar decays, independently of environmental context and of other ecological features of the total community.

The possibility that the assembly process is largely neutral apparently contradicts the observation that - locally - huge differences exist in plankton growth rates, leading to blooms visible even from space. However, the question here is whether, following the removal of the dominant species, such differences in growth rate are still important for explaining the abundance distribution. We could imagine that two species that bloomed in regions distant from the sampling location, and were transported by turbulence across more or less hostile environments, will have similar average demographic histories. As long as they find a niche they can thrive in, thus avoiding extinction, their abundance in other regions of the ocean should be largely independent of their specific adaptations. If the non-dominant part of the community is essentially composed of slow-growing organisms (for instance, in the form of resting stages), as hypothesized for the rare biosphere^10 9^, competitive ability in favourable conditions might be irrelevant. The similarity of the SADs power-law exponent to 3/2, a value obtained, for instance, in temporal spectra of intermittently varying ecosystems^45^, might indicate that local abundances bear a signature of spatio-temporal variability^27^.

All in all, understanding the interplay between ocean transport and community structure would require a resolution in space and time that is still unattained at a broad geographical scale. However, the combination of different observation platforms^46^, and the genomic characterization of samples in a given - usually coastal - location^3 47 48^ opens new perspectives for a dynamic characterization of drifting communities, which is not possible with the dataset we used.

Our interpretation of the observed ubiquitous trends in the plankton protist SADs, is based on features proper to the plankton lifestyle. It should thus not be expected that communities experiencing other ecological contexts and diverse assembly processes display the same regular pattern as evidenced in this work. Indeed, SADs reported for other microbial communities, in particular the gut microbiota, have been shown to obey different laws, possibly indicating different degrees of dispersal or experienced environmental variability. However, the current boost in molecular biodiversity surveys will undoubtedly help to clarify the universality of the observed pattern, and its likely relation with the peculiar interaction between ecology and physical forcing in the open ocean. Marine plankton populate 70% of Earth’s surface, providing the energy that fuels ocean food webs and contributing to global biogeochemical cycles. Understanding how these communities are assembled in their dynamic and highly variable environment, and the relation between rareness and biogeography^13 8 9 10 49^, paves the way to evaluating the resilience of marine biodiversity in a changing ocean.

## Methods

### Sequence data

Sequence data were retrieved from Richter et al.^30^. Briefly, seawater samples were collected in 121 stations distributed worldwide. Each sample was filtered using different mesh sizes so as to isolate different classes of the planktonic community: the pico-nanoplankton (0.8-5 μm), the nanoplankton (5-20 μm), the microplankton (20-180 μm), and the mesoplankton (180-2000 μm). Total DNA was extracted from each size class and the V9 region of the 18S rRNA gene^50^ was PCR-amplified and subjected to Illumina sequencing in order to describe eukaryotic planktonic communities. The retrieved paired-end sequencing reads were assembled using the fastx software (http://hannonlab.cshl.edu/fastx_toolkit/index.html) and dereplicated. To remove potential artefacts, we excluded sequences (i) that did not contain both forward and reverse primers, and (ii) identified as chimeras by the usearch programme (version 4.2; Edgar et al.^51^), using either the V9_PR2 reference database as reference (database W2; Guillou et al.^52^), or the *de novo* approach. The resulting sequences were clustered based on their similarity. First, the swarm algorithm^26^ was used to minimize the amount of PCR/sequencing errors, as well as intraspecific variability. Because we used it to cluster sequences distant from one mismatch, swarms represent the Operational Taxonomic Units (OTUs) at the finest taxonomic resolution. Second, swarms were clustered at 97% and 95% similarity using the UCLUST algorithm^53^ so as to obtain descriptions of the community structure at coarser taxonomic levels. All OTUs were then assigned to a taxon using the ggsearch program (Fasta package, http://faculty.virginia.edu/wrpearson/fasta/CURRENT/) against the V9_PR2 and the SILVA databases (http://www.arb-silva.de/projects/living-tree/; release LTPs111, February 2013). Finally, we excluded any swarm or OTU assigned to Bacteria, Archaea, Metazoa, Fungi or that were unassigned so as to focus only on the protist component of the dataset, thus allowing a consistent analysis of eukaryotic microbial communities in different samples.

### Species Abundance Distribution Model

Community assembly models allow to predict abundance distributions. Here we use the abundances of swarms or OTUs as a proxy for species abundances and do so in any particular sampling location and size class, from simple demographical hypothesis. Following He^38^, we assume that all *K* species in a local community undergo birth and death at rates that depend only on the species abundance (and not on its identity). For *n*≥1, the birth and death rates of a species of *n* individuals are:

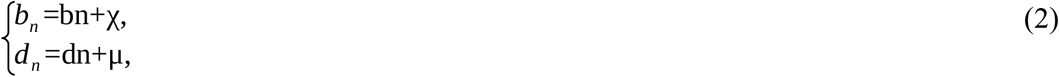

where *b* and *d* represent the linear per-capita birth and death rates while χ and μ are their non-linear corrections, whose effect is especially important for species of low abundance. When *n*=0, for consistency, *d*_*0*_=0 and *b*_*0*_=*v*+χ is the immigration/speciation rate (expressed as a sum for calculation convenience).

The solution of the master equation describing the ecological stochastic dynamics of such a modeled community provides the expected species abundance density distribution (*ϕ*_*n*_), that reads^54 19 38 39^:

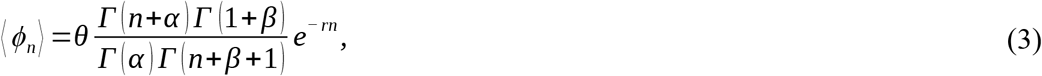

where:

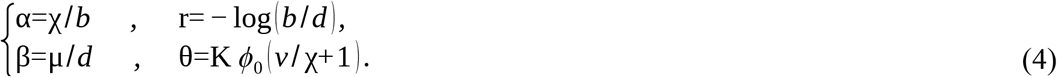

Note that the distribution, once normalized, depends on only three independent parameters (*r*,α,β); as in demographic terms linear birth and death rates compensate each other, so that only their relative value matters.

We found an asymptotic approximation (*n*>>1) of equation (3) (see Supplementary Information for the derivation) in the form:

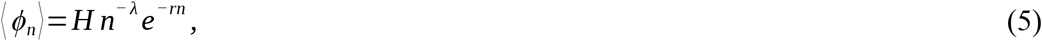

where *H* is a constant and λ=1-α+β. Equation (5) is an *exponentially truncated power-law*^20 55 56 57^ and for abundances of the order of a few tens of individuals and above, it approximates very well to the exact model distribution defined in equation (3). However, for low abundances, i.e., rare species, non-trivial combinations of the α and β parameters can cause detectable deviations from a pure power-law (see Fig. 1b). At first order, such deviations are controlled by a factor λ(α+β) (see Supplementary Information). Note finally that from equation (5) one can retrieve both the *exponential* (when α=β+1) and the *logseries* (when α=β) distribution in particular cases.

It is important to stress that for the general case in which the non-linear terms account for both dispersal and density dependent effects, their respective contributions to equations (2) can not be disentangled. Therefore, no univocal conclusion can be drawn about the relative effect of local ecological dynamics versus dispersal from fitting the model to the observational data. Nevertheless we can consider that, in the oceanic environment, dispersal has related effects on both *d*_*n*_ and *b*_*n*_. Indeed, the demographic parameters at the local scale are affected by the water flux through the boundary of the (supposedly well-mixed) sampling region. Because the movement of oceanic surface waters can be considered as an approximately incompressible planar flow, the inflow equals the outflow. For planktonic species whose abundance is sufficiently large (for which density-dependent terms can be neglected) and uniform at a scale bigger than that where the local community is sampled, the effect of transport-induced dispersal on the demography can be represented by a same rate *t*. In a linear approximation, we can write *b*=*b̅*+*t* and *d*=*d̅*+*t*, where the coefficients *b̅* and *d̅* are the per capita local rates of cells division and death. If dispersal dominates local population dynamics, hence *t*≫*b̅*,*d̅*, then the community will be effectively close to equilibrium. Such situation would result in a vanishing exponential cutoff (i.e. r~0) and, correspondingly, in an extended power-law regime for the Species Abundance Distribution (equations (3) and (5)).

### Model fitting and testing

We developed an algorithm to adaptively fit the theoretical model to the empirical distributions, which allows us to automatically distinguish between two components of the community: a part that is described by the model within a fixed confidence level, and a part of dominant species whose abundance cannot be described by the same model within the same confidence level. We fitted the data by maximum likelihood. Indeed, alternative methods - such as least-squares minimization - can produce substantially inaccurate estimates^36 58^. To assess the significance of the obtained fits we rely on p-value calculation and not on the fraction r^2^ of variance accounted for by the fits, as the latter has been criticized as having little power as a hypothesis test indicator^36 59^. We applied the same adaptive algorithm to the previously introduced model (equation (3)) and to other theoretical models commonly used in SAD model selection studies: the exponential, log-series, log-normal, Poisson log-normal and zero-sum multinomial distributions.

The fits, whose parameters are summarized in Table 1, are obtained for each sample by iteration of four steps: 1) maximize the likelihood of the distribution of a subset of species whose abundance is smaller than a given threshold, thus obtaining the best fitting parameters; 2) compute a measure of the distance between the fit and the data; 3) evaluate, by bootstrapping, the statistical significance of the fit and generate numerical p-values; 4) repeat the above steps for increasing values of the abundance threshold and select the largest subset of species such that the p-value associated with its fit is larger than a given confidence.

Observed abundances of swarms/OTUs in a specific sample, from now on denoted for simplicity ‘species’, are arranged in order of increasing abundance in the vector ***n***=(*n*_*1*_, … *n*_*k*_ …, *n*_*K*_), such that *n*_k_ ≤ *n*_*k*+*1*_. Hence *n*_*k*_ is the number of individuals (abundance) of species k, *N* = Σ *n*_*k*_ is the total number of individuals, and *K* is the number of species observed in the local community. The typical representation of such observations is obtained by counting the number of species whose abundance belongs to fixed intervals of abundances (the empirical SAD). Fig. 1b represents such a histogram in logarithmic binning, that is most adapted for visualizing the SAD in double logarithmic scale. In our study, we focus on subset of 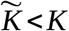 species, whose abundances 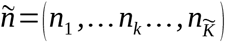 are smaller than a cutoff 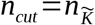, which we denote sub-community. For statistical purposes, we are interested in the associated cumulative distribution function 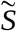 defined as:

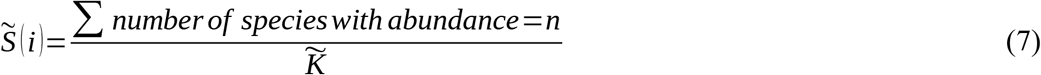

The *discrete upper-bounded model distribution p(n)* is defined from equation (3), by imposing an upper limit to the abundances, as:

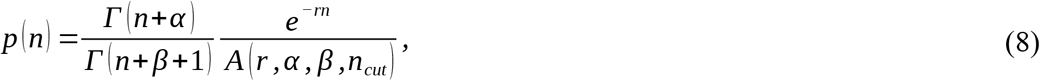

where *A*(*r*,α,β,*n*_cut_) is the normalization constant. In the limit of *n*_*max*_→ ∞, we find *p*(*n*) → (*ϕ*_*n*_). The correspondent cumulative distribution is denoted as *P*(*i*)=Σ *P*(*n*).

Step 1 of the adaptive loop allows to obtain the parameters for the best fit to any given sub-community data by *likelihood maximization*. We define the log-likelihood of the data given the model distribution in equation (8) as:

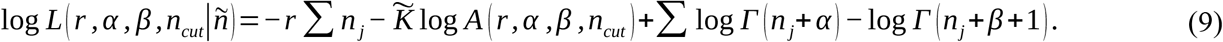

Equation (9) is maximized using a *generalized simulated annealing* algorithm^60^ to retrieve the best estimates of the model parameters (*r*,α,β) for a fixed *n*_*cut*_. This approach improves the optimization process by minimizing the risk of remaining trapped in a local minimum.

In order to evaluate the goodness of the fit, we need to introduce a metric of the distance between the empirical and the theoretical distributions (step 2). To this aim, we chose the *Anderson-Darling distance*^61^, best adapted to detect differences in low-abundance classes than the more commonly used Kolmogorov-Smirnov statistics. The Anderson-Darling metric D between the cumulative distributions reads:

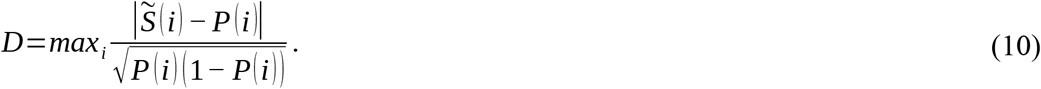

The distance between distributions, however, does not provide the probability that the observed data are obtained as a random sample from the theoretical distribution. Indeed, the intrinsically discrete structure of the distributions does not allow the use of critical values tables for statistical tests, so that a p-value cannot be directly assigned from the Anderson-Darling test.

We adopt thus the bootstrapping approach followed by Clauset et al.^36^ (step 3). The rationale is that that best fits to the data obtained in step 1 shall be discarded if their distance from the empirical data is unlikely due to random fluctuations. Such random deviations are quantified by the statistics of the distances between random samples of the best fitting model distribution and their relative maximum likelihood fits. For each fit, we generated 200 synthetic datasets, whose 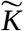 abundances match the length of the vector ñ, by sampling the model distribution *p*(*n*) for the best fitting parameters (*r*,α,β). We then measure the Anderson-Darling distance of each such synthetic dataset from the distribution that fits them with highest likelihood. Finally, we assign a *p-value* to the original fit as the fraction of those distances that are larger than the Anderson-Darling distance between the original data and its best fitting distribution. Hence, fits are considered significant if the empirical distance falls within ‘typical’ values that we would expect to obtain by random sampling. In this case, the fraction p is large. Instead, p is small if the samples are systematically less distant from their fit than the empirical distribution is from its own fit. By setting a minimum threshold on p (in our analysis, 0.1), we aim to exclude the latter case.

We repeated the maximum-likelihood fit and p-value calculation for sub-communities of growing size (step 4), by progressively increasing *n*_*cut*_ (starting from *n*_*cut*_ = 100). We select then the largest sub-community compatible with the p-value threshold and we call *n*_*max*_ its associated *n*_*cut*_. The p-value and *n*_*max*_ of this sub-community is displayed for every sample in Supplementary Data 1. Our method allows the identification of *n*_*max*_ even when the Anderson-Darling statistics do not present a marked minimum in such interval. However, it is computationally more intensive than the method proposed by Clauset et al.^36^, where the p-value was only computed for the sub-community that minimizes the Kolmogorov-Smirnov distance. For this reason, the adaptive fit was bounded to a maximum abundance *n*_*cut*_ of 10100, and still numerical calculations were only possible on a computer cluster. Correspondingly, the extension of the fits reported in Table 1 underestimate the real maximal abundance that could be accommodated by the model, thus the size of the non-dominant component of the community.

In the adaptive algorithm, the level of confidence for statistical significance affects the range of abundances that fall within the non-dominant component of the community. If the confidence level is too high (so that the statistical test is too stringent), the fit will not accommodate for the inevitable noise induced by measurement errors, fitting will be poor for any theoretical distribution, and the identification of the non-dominant component inefficient. If it is too low, the adaptive fit will always succeed, and the whole distribution will likely be described by one single law, whose statistical significance will however be poor. We chose a confidence level of p=0.1, which is more stringent than what was used in previous analyses. We checked that small changes in the confidence level did not affect the results: relaxing the tolerance by decreasing the threshold p-value to 0.05 led to a few more stations to be fitted by equation (3), and slightly larger non-dominant components, but only a negligible enhancement of the variability in the distributions of the power-law exponent.

## Acknowledgements

The authors are very grateful to Francesco d’Ovidio, Valentin Anjou and Stéphane Audic, who participated in the early stages of this work, to Olivier Missa, Guilhem Sommeria-Klein and Erik van Nimwegen for discussions on neutral models and model fitting, to the *Tara* Oceans consortium, and to three anonymous referees for their constructive comments. The support of the informatics platform of IBENS has been essential for the computational part of this study. This work has received support under the programmes «Investissements d’Avenir» launched by the French Government and implemented by ANR with the references ANR-10-LABX-54 MEMOLIFE, ANR-10-IDEX-0001-02 PSL* Research University and OCEANOMICS, as well as from the EU project MicroB3. CB additionally acknowledges funding from the ERC Advanced Award “Diatomite”, the Louis D Foundation of the Institut de France, and the Radcliffe Institute of Advanced Study at Harvard University for a scholars fellowship during the 2016-2017 academic year. This research was supported in part by the National Science Foundation Grant No. NSF PHY-1125915, NIH Grant No. R25GM067110, and the Gordon and Betty Moore Foundation Grant No. 2919.01. This article is contribution number XX of *Tara* Oceans.

## Author Contributions

S.D.M. conceived and directed the study. E.S-G. obtained the analytical results, designed the adaptive algorithm and produced the fits. S.M. carried out the preliminary analysis on protist RADs. L.Z. performed statistical analysis on fitted parameters. S.D.M., E.S-G., L.Z. and C.B. interpreted the results and wrote the manuscript. C.d.V. and E. K. provided access to the *Tara* Oceans dataset and commented on the manuscript.

## Author Information

The authors declare no competing financial interests. Correspondence and requests for materials should be addressed to Silvia De Monte.

## References

1. Sunagawa, S. et al. Structure and function of the global ocean microbiome. Science 348, 1261359 (2015).

2. Vargas, C. de et al. Eukaryotic plankton diversity in the sunlit ocean. Science 348, 1261605 (2015).

3. Fuhrman, J. A. Microbial community structure and its functional implications. Nature 459, 193–199 (2009).

4. Hutchinson, G. E. The Paradox of the Plankton. Am. Nat. 95, 137–145 (1961).

5. d’Ovidio, F., Monte, S. D., Alvain, S., Dandonneau, Y. & Lèvy, M. Fluid dynamical niches of phytoplankton types. Proc. Natl. Acad. Sci. 107, 18366–18370 (2010).

6. Hanson, C. A., Fuhrman, J. A., Horner-Devine, M. C. & Martiny, J. B. H. Beyond biogeographic patterns: processes shaping the microbial landscape. Nat. Rev. Microbiol. 10, 497–506 (2012).

7. McGillicuddy, D. J. Mechanisms of Physical-Biological-Biogeochemical Interaction at the Oceanic Mesoscale. Annu. Rev. Mar. Sci. 8, 125–159 (2016).

8. Galand, P. E., Casamayor, E. O., Kirchman, D. L. & Lovejoy, C. Ecology of the rare microbial biosphere of the Arctic Ocean. Proc. Natl. Acad. Sci. 106, 22427–22432 (2009).

9. Pedrós-Alió, C. The Rare Bacterial Biosphere. Annu. Rev. Mar. Sci. 4, 449–466 (2012).

10. Lennon, J. T. & Jones, S. E. Microbial seed banks: the ecological and evolutionary implications of dormancy. Nat. Rev. Microbiol. 9, 119–130 (2011).

11. Lynch, M. D. J. & Neufeld, J. D. Ecology and exploration of the rare biosphere. Nat. Rev. Microbiol. 13, 217–229 (2015).

12. Sogin, Mitchell L., et al. Microbial diversity in the deep sea and the underexplored “rare biosphere”. Proceedings of the National Academy of Sciences 103(32), 12115–12120 (2006).

13. Logares, R. et al. Patterns of Rare and Abundant Marine Microbial Eukaryotes. Curr. Biol. 24, 813–821 (2014).

14. Barberán, A., Casamayor, E. O. & Fierer, N. The microbial contribution to macroecology. Front. Microbiol. 5, (2014).

15. Chust, G., Irigoien, X., Chave, J. & Harris, R. P. Latitudinal phytoplankton distribution and the neutral theory of biodiversity. Glob. Ecol. Biogeogr. 22, 531–543 (2013).

16. Locey, K. J. & Lennon, J. T. Scaling laws predict global microbial diversity. Proc. Natl. Acad. Sci. 113, 5970–5975 (2016).

17. McGill, B. J. et al. Species abundance distributions: moving beyond single prediction theories to integration within an ecological framework. Ecol. Lett. 10, 995–1015 (2007).

18. Magurran, A. E. & McGill, B. J. Biological Diversity: Frontiers in Measurement and Assessment. (Oxford University Press, 2011).

19. Volkov, I., Banavar, J. R., Hubbell, S. P. & Maritan, A. Neutral theory and relative species abundance in ecology. Nature 424, 1035–1037 (2003).

20. Pueyo, S. Diversity: between neutrality and structure. Oikos 112, 392–405 (2006).

21. Connolly, S. R. et al. Commonness and rarity in the marine biosphere. Proc. Natl. Acad. Sci. 111, 8524–8529 (2014).

22. Ulrich, W., Ollik, M. & Ugland, K. I. A meta-analysis of species-abundance distributions. Oikos 119, 1149–1155 (2009).

23. Shoemaker, W. R., Locey, K. J. & Lennon, J. T. A macroecological theory of microbial biodiversity. Nature Ecology & Evolution 1, 0107 (2017).

24. Matthews, T. J. & Whittaker, R. J. Neutral theory and the species abundance distribution: recent developments and prospects for unifying niche and neutral perspectives. Ecol. Evol. 4, 2263–2277 (2014).

25. Baldridge, Elita, et al. An extensive comparison of species-abundance distribution models. PeerJ 4, e2823 (2016).

26. Xiao, Xiao, James P. O’Dwyer, and Ethan P. White. Comparing process-based and constraint-based approaches for modeling macroecological patterns. Ecology 97(5), 1228–1238 (2016).

27. Azaele, S., Pigolotti, S., Banavar, J. R., & Maritan, A. Dynamical evolution of ecosystems. Nature 444, 926–928 (2006).

28. Magurran, Anne E., and Peter A. Henderson. Explaining the excess of rare species in natural species abundance distributions. Nature 422, 714–716 (2003).

29. Ulrich, W., & Ollik, M.. Frequent and occasional species and the shape of relative-abundance distributions. Diversity and distributions, 10(4), 263–269 (2004).

30. Richter et al. Global plankton biogeography is shaped via ocean circulation dynamics. Under review (2017).

31. Mahé, F., Rognes, T., Quince, C., Vargas, C. de & Dunthorn, M. Swarm v2: highly-scalable and high-resolution amplicon clustering. PeerJ 3, e1420 (2015).

32. Brown, J. H., Gillooly, J. F., Allen, A. P., Savage, V. M. & West, G. B. Toward a Metabolic Theory of Ecology. Ecology 85, 1771–1789 (2004).

33. Woodward, G. et al. Body size in ecological networks. Trends Ecol. Evol. 20, 402–409 (2005).

34. White, E. P., Ernest, S. K. M., Kerkhoff, A. J. & Enquist, B. J. Relationships between body size and abundance in ecology. Trends Ecol. Evol. 22, 323–330 (2007).

35. Matthews, T. J., Borges, P. A., Azevedo, E. B., & Whittaker, R. J. A biogeographical perspective on species abundance distributions: recent advances and opportunities for future research. Journal of Biogeography, 44(8), 1705–1710 (2017).

36. Clauset, A., Shalizi, C. & Newman, M. Power-Law Distributions in Empirical Data. SIAM Rev. 51, 661–703 (2009).

37. Jeraldo, P. et al. Quantification of the relative roles of niche and neutral processes in structuring gastrointestinal microbiomes. Proc. Natl. Acad. Sci. 109, 9692–9698 (2012).

38. He, F. Deriving a neutral model of species abundance from fundamental mechanisms of population dynamics. Funct. Ecol. 19, 187–193 (2005).

39. Azaele, S. et al. Statistical mechanics of ecological systems: Neutral theory and beyond. Rev. Mod. Phys. 88, 035003 (2016).

40. Volkov, I., Banavar, J. R., He, F., Hubbell, S. P. & Maritan, A. Density dependence explains tree species abundance and diversity in tropical forests. Nature 438, 658–661 (2005).

41. Chaffron, S. et al. Environmental context of selected samples from the Tara Oceans Expedition (2009-2013). Pangaea, (2014). doi:doi:10.1594/PANGAEA.840718

42. Tittensor, D. P. et al. Global patterns and predictors of marine biodiversity across taxa. Nature 466, 1098–1101 (2010).

43. Gravel, D., Canham, C. D., Beaudet, M. & Messier, C. Reconciling niche and neutrality: the continuum hypothesis. Ecol. Lett. 9, 399–409 (2006).

44. Wilkins, D., Van Sebille, E., Rintoul, S. R., Lauro, F. M., & Cavicchioli, R.. Advection shapes Southern Ocean microbial assemblages independent of distance and environment effects. Nature communications, 4, 2457 (2013).

45. Ferriere, R. & Cazelles, B. Universal Power Laws Govern Intermittent Rarity in Communities of Interacting Species. Ecology 80, 1505–1521 (1999).

46. Lehahn, Y., d’Ovidio, F., & Koren, I.. A Satellite-Based Lagrangian View on Phytoplankton Dynamics. Available online ahead of print Annual review of marine science (2017).

47. Cram, J. A., Chow, C. E. T., Sachdeva, R., Needham, D. M., Parada, A. E., Steele, J. A., & Fuhrman, J. A. Seasonal and interannual variability of the marine bacterioplankton community throughout the water column over ten years. The ISME journal, 9(3), 563 (2015).

48. Martin-Platero, A. M., Cleary, B., Kauffman, K., Preheim, S. P., McGillicuddy, D. J., Alm, E. J., & Polz, M. F.. High resolution time series reveals cohesive but short-lived communities in coastal plankton. Nature communications, 9(1), 266 (2018).

49. Magurran, A. E. Measuring biological diversity. Blackwells. (Oxford, UK, 2004).

50. Amaral-Zettler, L. A., McCliment, E. A., Ducklow, H. W. & Huse, S. M. A Method for Studying Protistan Diversity Using Massively Parallel Sequencing of V9 Hypervariable Regions of Small-Subunit Ribosomal RNA Genes. PLOS ONE 4, e6372 (2009).

51. Edgar, R. C. UPARSE: highly accurate OTU sequences from microbial amplicon reads. Nat. Methods 10, 996–998 (2013).

52. Guillou, L. et al. The Protist Ribosomal Reference database (PR2): a catalog of unicellular eukaryote Small Sub-Unit rRNA sequences with curated taxonomy. Nucleic Acids Res. 41, D597–D604 (2013).

53. Edgar, R. C. Search and clustering orders of magnitude faster than BLAST. Bioinformatics 26, 2460–2461 (2010).

54. McKane, A., Alonso, D. & Solé, R. V. Mean-field stochastic theory for species-rich assembled communities. Phys. Rev. E 62, 8466–8484 (2000).

55. Dennis, B. & Patil, G. P. The gamma distribution and weighted multimodal gamma distributions as models of population abundance. Math. Biosci. 68, 187–212 (1984).

56. Pueyo, S., He, F. & Zillio, T. The maximum entropy formalism and the idiosyncratic theory of biodiversity. Ecol. Lett. 10, 1017–1028 (2007).

57. Green, J. L. & Plotkin, J. B. A statistical theory for sampling species abundances. Ecol. Lett. 10, 1037–1045 (2007).

58. Goldstein, M. L., Morris, S. A. & Yen, G. G. Problems with fitting to the power-law distribution. Eur. Phys. J. B - Condens. Matter Complex Syst. 41, 255–258 (2004).

59. Bauke, H. & others. Parameter estimation for power-law distributions by maximum likelihood methods. Eur. Phys. J. B Condens. MATTER Phys. 58, 167 (2007).

60. Tsallis, C., & Stariolo, D. A. Generalized simulated annealing. Physica A: Statistical Mechanics and its Applications, 233(1-2), 395–406 (1996).

61. Anderson, T. W., & Darling, D. A. Asymptotic theory of certain” goodness of fit” criteria based on stochastic processes. The annals of mathematical statistics, 193–212 (1952).

